# Methodology for inference of intercellular gene interactions

**DOI:** 10.1101/2023.02.26.530111

**Authors:** Saurabh Modi, Ryan Zurakowski, Jason P. Gleghorn

## Abstract

To govern organ size, shape, and function, cell-secreted diffusible molecules called morphogens spatially pattern cell differentiation, gene expression, and proliferation. Local morphogen concentration governs cell differentiation through gene regulatory networks (GRN). Previous inference methodologies tackle intercellular GRN inference between cells of one type. This is insufficient, as many developmental systems consist of cells of different types interacting with each other. Inference methodologies of GRNs between different cell types assume knowledge of diffusible morphogen identity and concentration. This makes their applicability limited in real biological systems. Here, we develop a computational methodology to infer the intercellular GRN derived from experiments that use fluorescence from reporter proteins for gene expression measurements. For validation, we demonstrate the methodology *in silico* using three case studies based on developmental and synthetic biology. The results show that, barring practical identifiability limitations, the methodology successfully infers the intercellular GRNs.

## 1 Introduction

In developing animals and plants, extracellular diffusible molecules called morphogens are secreted from cells that coordinate spatial patterning of cell differentiation, gene expression, and proliferation to regulate organ size, shape, and function [1–8]. For example, sonic hedgehog is secreted by mesoderm cells on the ventral side of the spinal cord in vertebrates. This generates a morphogen concentration gradient throughout the developing spinal cord and defines differentiation events encoded by intracellular gene regulatory networks (GRN) [9, 10]. This multicellular interaction is an example of sender-receiver type signaling, where morphogen secretion from one cell type induces a response from another cell type. Often deviations in the strength or positioning of morphogen concentrations and gradients regulate this intercellular interaction and underpin developmental diseases and congenital birth defects [11–16]. Since these interactions function at the multicellular level (tissue scale) with spatial heterogeneity, many traditional methods of inferring the gene regulatory networks that comprise these interactions are insufficient.

Single-cell sequencing technology has significantly advanced the inference of underlying GRNs [17–21]. Complete single-cell RNA-seq (scRNA-seq) datasets can be correlated to spatial positions by mapping to landmark gene *in situ* hybridization spatial maps resulting in connections between genes interacting across cells based on their mutual proximity [22–25]. However, regulatory connections and rate parameters encoded in dynamic information are not possible to infer as this methodology only provides data at a single snapshot in time [26]. Previous analyses that tackle the inference of nonlinear mechanistic GRNs do not consider multiple interacting cell types that may have different intracellular GRNs [27, 28]. This is a common feature of multicellular developmental systems where two different cell types may interact to give different shapes and sizes of organs and organisms [29–34].

This work develops a systematic methodology to assay the intercellular GRN and a computational framework to use the generated data to infer its connections and parameters. Studies quantifying the gene interaction between two cell types often involve the construction of dose response curves to obtain the Hill parameters of interaction [35, 36]. However, morphogens that mediate intercellular interactions are often not fully known, known, and the degradation rate and morphogen diffusion constants are often not addressed. These parameters are critically important for interactions over large distances where morphogen gradients are relevant. This is the first study with a methodology to infer intercellular GRN parameters agnostic of the morphogen concentration measurement. In place of morphogen quantification, our method uses fluorescence measurements from reporter proteins to infer intercellular GRNs. These proteins are expressed from reporter genes, placed under the regulation of the same promoter as the gene of interest [37]. In addition, we propose simple longitudinal experiments and 2D spatial co-culture imaging experiments to generate the necessary spatiotemporal data required for the inference. Whereas we lose the ability to infer the absolute values of the interaction parameters with concentration units, we can infer their scaled versions and diffusion constants. To determine if the resulting model is identifiable, we also introduce a framework for identifiability analysis of the model with scaled parameters given spatiotemporal reporter expression data.

## 2 Methodology for inference

### 2.1 Reaction-diffusion intercellular interaction framework

To infer the interaction parameters, we needed a computational framework incorporating diffusible factor secretion, diffusion, and its interplay with intracellular gene expression. Many approaches have been proposed to model cell-cell interactions based on diffusion and reaction equations [29, 38, 39]. We adopted a similar 2D spatiotemporal system described by the partial differential equation:

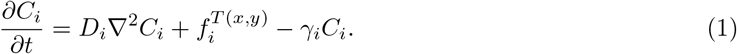

Here *C*_*i*_ is the concentration of the *i*^*th*^ species and *D*_*i*_ is its diffusion coefficient through the medium. We assume species are secreted from sender cells by passive diffusive transport across a permeable cell membrane. The morphogen binds to receptors on the receiver cell membrane and initiates intracellular gene expression mediated by a signaling cascade. The binding and unbinding to receptors at the cellular level sequesters the morphogen and affects its bulk diffusion through the extracellular medium. These processes can be lumped into a bulk diffusion constant *D*_*i*_, and consequently, we neglect the corresponding detailed transport equations [40]. We assume that the extracellular medium is homogeneous, resulting in a constant *D*_*i*_. For intracellular species that do not diffuse out of the cell, the diffusion term is absent (*D*_*i*_ = 0). This includes reporter proteins, making passive diffusion through cell membranes difficult due to their large molecular weights. Each species is produced from its gene with the rate *f*_*i*_ and degrades intracellularly at a rate *γ*_*i*_. The extracellular morphogen degradation rate is assumed to be negligible compared to the corresponding intracellular rate. This is because active degradation processes like proteolysis (in sender cells) or receptor-mediated degradation (in receivers) occur at a higher rate than uncatalyzed extracellular protein decay. Further, the degradation rate of the morphogen is assumed to be the same across all cells (senders and receivers). To allow for gene regulation, we let the production rate *f*_*i*_ = *f*_*i*_(*C*_1_, *C*_2_…*C*_*r*_) be a function of the concentration of other species that act as transcription factors. Reporter proteins have the same production rate *f*_*i*_ as their corresponding gene of interest as they share a promoter. Local extracellular morphogen concentration around receiver cells is assumed to regulate intracellular gene expression through the function *f*_*i*_ even though the mechanism of action is indirect, i.e., binding to membrane receptors and inducing a signaling cascade with downstream gene regulation. This assumption is supported by the fact that in typical mammalian systems, the signaling cascade responds at a much faster time scale compared to that of gene expression [41, 42]. Each species is assumed to be produced from one cell type; hence the production of the *i*^*th*^ species 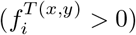 occurs only at the coordinates (*x, y*) where the cells of the corresponding type *T* (*x, y*) exist.

The function *f*_*i*_ can be an increasing (or decreasing) function with respect to its argument in the case of a transcriptional activator (repressor). In general, *f*_*i*_ can take the form of a Hill function with respect to transcription factors. Further, multiple transcription factors acting on the same gene can act as sufficient, necessary, or a combination of these [27]. We used the multiplication operator for the necessary condition and summation for the sufficient condition. We illustrate this with an example where the gene expression is repressed by *C*_1_ and activated by *C*_2_ as necessary regulation and activated by *C*_3_ and *C*_4_ as sufficient regulation. Then the resulting production rate can be written as:

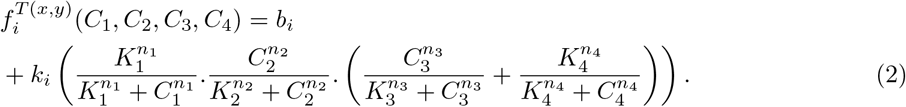

To solve the PDE in (1), we numerically simulated the system, where a region within a cell culture dish was discretized into uniform control volumes of size *δx* (**Figure 1**). Each control volume was small enough to overcome diffusion limitations and thus was assumed to be spatially homogeneous. Diffusion between control volumes was based on the flux of a species through the face of the control volume in accordance with the finite volume method [43]. For simplicity, we assumed that the cells were distributed spatially in a 2D geometry with defined boundaries between cell types. Each control volume consisted of only one type of cell or was empty. A no flux boundary condition was used for diffusible factors consistent with a closed system like the petri dish.

**Figure 1.**
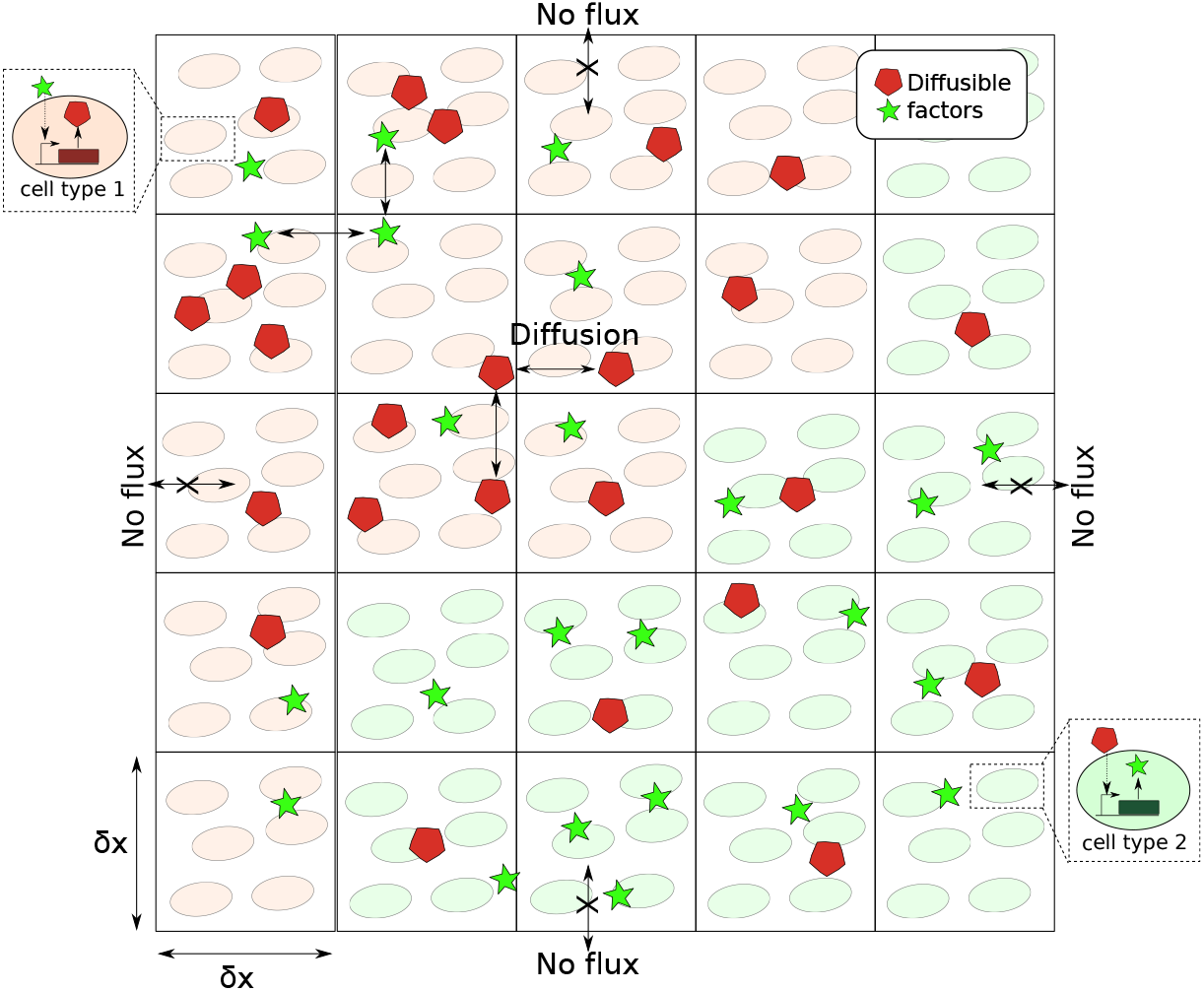
A schematic representation of the intercellular gene interaction framework with morphogen diffusion and gene expression in an example system, simulated using the finite volume method.

### 2.2 Rate parameter identifiability

Before the task of parameter inference, identifiability analysis was used to determine the parameters or combinations that were uniquely identifiable given a particular set of measured outputs and initial conditions. Accordingly, inferred parameters were determined or constrained based on their combinations.

It is important to note that we took the measurement variable as the receiver reporter-protein concentration in terms of its fluorescence measurement. The question of how to use the arbitrary linear scaling of concentration (fluorescence intensity) to give parameters was addressed by normalizing the fluorescence with its maximal spatiotemporal value (discussed later). To demonstrate the scaling of parameters when the receiver reporter fluorescence was available, we started with the differential equations describing the spatiotemporal sender-receiver type interaction of gene *P* (diffusible gene product) and *Q* (reporter):

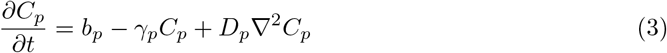

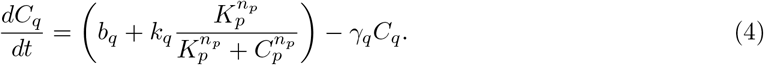

Here the Hill function represents the transcriptional regulation of *P* on *Q*, with only *C*_*q*_ being the measured state. The parameters *b*_*p*_ and *K*_*p*_ can be simultaneously scaled up or down without affecting *C*_*q*_. Additionally, as the reporter measurement (*C*_*q*_) is in arbitrary units, the production parameters of *Q*, namely *b*_*q*_ and *k*_*q*_, are not uniquely identifiable. Hence we re-scale the parameters as follows:

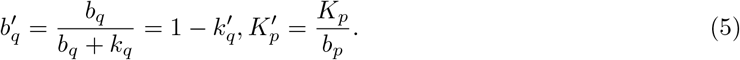

Here 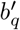 and 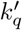 are the proportion of the total production rate contributed by basal expression and the proportion which is repressible by *P*, respectively. The amount of time required to accumulate enough *P* (starting from zero) to affect half the maximum repression of *Q* is given by 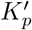. Next, an identifiability analysis algorithm was executed after re-scaling parameters according to the above scheme for all such regulator and regulated species pairs in an intercellular GRN to determine which re-scaled parameters were uniquely identifiable.

We used the package STRIKE-GOLDD [44] to implement the identifiability analysis. We took the measured output to be the spatiotemporal fluorescence of the reporter protein in the two cell types. We then built the mathematical model to simulate the spatiotemporal evolution of the measured variable (reporter fluorescence). Meshing 2D space with many control volumes accurately captures the fluorescence gradient but leads to excessive computational expense. Instead, we considered a 3 *×* 3 configuration with no flux boundary conditions as the simplest geometry that could still express fluorescence gradients at a low resolution. If we show that parameters are identifiable in this simple geometry, they will be identifiable with a larger number of *n × n* control volumes, as this only serves to increase spatial resolution. The same assumptions from the reaction-diffusion framework were applied here also. The control volumes were asymmetrically assigned cell types, so that four were of one cell type and five of the other cell type. This avoided redundant fluorescence measurements due to equidistant sender cell and receiver cell control volumes.

The differential equations for each element were then built using the Symbolic Math Toolbox in MATLAB based on the corresponding cell type’s internal GRN and neighboring elements with which diffusible factors are exchanged. We then applied the STRIKE-GOLDD algorithm to this ODE system that described the spatiotemporal evolution of measured reporter gene expression. The initial conditions for each cell type were set as the steady-state value of variables without intercellular interactions (diffusion constants=0). In this way, the initial condition was a function of the parameters. We also assume in this analysis that the Hill coefficient is equal to 1 as in the general case (*>* 0), a symbolic solution for the initial condition was intractable. In some instances, investigating the entire model for identifiability is computationally inefficient; hence we opted for numerical analysis and model decomposition. Whereas this analysis does not rigorously address the question of the identifiability of 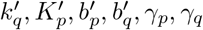, and *D*_*p*_ given the Hill coefficient *n*_*p*_ as a free variable to be estimated, the answer should not be different from the identifiability of the parameters given *n*_*p*_ = 1. Intuitively, *n*_*p*_ is a dimensionless parameter independent of the linear scaling of *K*_*p*_ and the regulator species concentration, which is proportional to the simultaneous scaling of *b*_*p*_. In other words, in the toy model, the combinations *b*_*q*_*/*(*b*_*q*_ + *k*_*q*_) and *K*_*p*_*/b*_*p*_ are identifiable independent of a fixed value of *n*_*p*_. The Hill coefficient is identifiable as it represents the steepness of the scale-invariant spatial gradient of the inducible reporter protein fluorescence measurement. The application of this identifiability analysis to support inference in case studies will be shown in section 3.

### 2.3 Experimental data needed for inference

Let there be two cell types with genes of interest, each (with corresponding reporter proteins) interacting through the intercellular GRN. We propose two experiments to assay the GRN and generate spatiotemporal reporter expression data. These data can then be used in our computational framework to infer the connections and parameters in the GRN.

1. Perform a conditioned medium experiment by culturing cells separately and using the collected conditioned medium from one cell type as the growth medium for the other cell type. Measure the fluorescence in a plate reader with respect to time and normalize to the cell number using the optical density of the cell suspension. After sufficient induction time, replace the used medium with fresh medium and obtain the time profile of this expression. It is important to note that the resulting fluorescence data have only time as the independent variable, with any spatial heterogeneity being averaged out. This experiment determines the internal dynamics of each cell type including the degradation rate of reporter proteins of the gene of interest. Following this experiment, repeat using the collected conditioned medium from the other cell type to query for reciprocal signaling.
2. Co-culture cells within defined spatial zones to allow for image-based fluorescent quantification of spatial 2D gene expression patterns. The gene expression of reporter proteins in the cells changes with time and is imaged longitudinally at regular time intervals. In these experiments, care must be taken to limit the growth rate of cells to avoid oversaturation of fluorescence during live imaging. The basal per-cell fluorescence expression in growth-reduced cells must be equal to the controls. This ensures that transcriptional and translational processes in regulating the genes of interest are not heavily affected by growth suppression.

### 2.4 A computational framework for intercellular GRN inference

The computational inference framework was based on model fitting to experimental data. First, a genetic algorithm inferred the types of interactions between genes including activation, repression, or no interaction. Then given the structure of the model and its equations, continuous parameters such as the rate parameters and diffusion constants were inferred using a direct-search optimization algorithm.

#### 2.4.1 Genetic algorithm for edge type inference

The goal of the genetic algorithm (GA) was to infer the smallest network of genes (nodes) and the type of interactions (activation/repression/none) defined by directed edges between them that described the experimental data. Even though edge type inference with discrete variables is the objective of the GA, continuous parameters need to be incorporated as they are part of the functional relationship between regulating and regulated genes. The genetic algorithm is ideal for this task as it allows for a combination of discrete and continuous parameters similar to the method stipulated in [27]. Further, it handles discrete network edge variables like type of interaction differently than continuous variables such as the Hill parameters. The fine-tuned inference of continuous parameters was carried out after convergence of the edge types, using algorithms specialized for such a task (described in the next section). This improved computational efficiency as the GA would continually query different edge types and continuous parameter values after network edge type convergence. For a set of network parameters, *in silico* data simulating experiments in section 2.3, were generated using the computational framework described in section 2.1. A measure of fit was needed to fit the simulations to the experimental data. To this end, we used an Akaike-like information criterion given by:

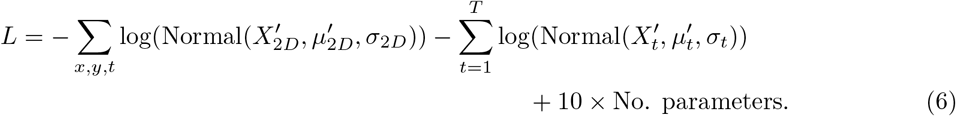

The first two terms on the right-hand side are the log-likelihood function from the 2D co-culture and the conditioned media experiment, respectively. This was used to quantify the probability that a particular network configuration explains the experimental measurements. A Gaussian probability function was chosen in the likelihood function as a heuristic approach to reproduce the spatiotemporal profile without bias in the weightage of data points based on their absolute values. The scaled measurements and simulated values were defined as:

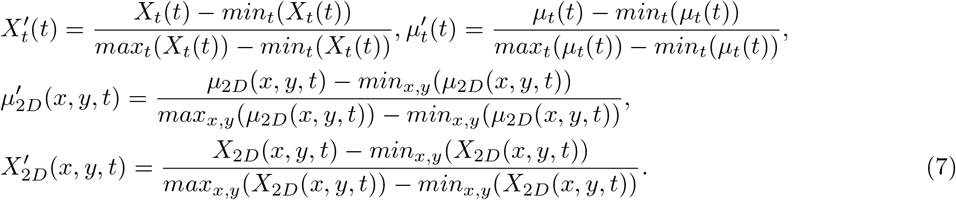

Here *X*_*t*_ and *X*_2*D*_ represent the fluorescence measurement from the conditioned media experiment and the 2D co-culture experiment, respectively, while *µ*_*t*_ and *µ*_2*D*_ are the corresponding simulated values. The fluorescence measurements from the 2D co-culture experiment are spatially averaged within each control volume boundary. The standard deviation parameters *σ*_*t*_ and *σ*_2*D*_ represent the measurement noise of the respective experiments. The data scaling between the minimum and maximum values are necessary to improve edge type inference by amplifying the spatiotemporal dynamics of cells. The shape of the response is critical to identifying the nature of the extracellular induction and intracellular gene regulation. To reduce noise, the temporal gene expression data were transformed using a five-point local moving average before scaling according to (7). Further, uninformative data was removed by truncating the time span for simulated and experimental data to the time point *T*, where the temporal expression was relatively steady. The linear scaling of fluorescence measurement between the minimum and maximum values amplifies the nature of the input step response of the cells to inducing factors. The final term in (6) penalizes overfitting the data by unnecessary network edges. A higher penalty multiplier (*×*10) is used compared to the Akaike criterion to accelerate the convergence of networks and reduce computational time for each run. Note that the objective of this GA was to infer a network that minimizes the value *L*.

With the scoring system defined to quantify the goodness of fit of the experimental data to the network configuration, we were able to search and reverse engineer the intra/intercellular interaction networks. The genetic algorithm works by maintaining a population of network configurations that evolve in parallel for a given number of generations. To maintain the diversity of networks, we initialized the first generation with a population of random edge-dense networks. Then the following steps were executed sequentially within each generation:

1. Generate a population of candidate networks (children) using ones from the previous generation by using two operations: mutation and crossover. Two parents of the prior generation were randomly chosen to carry out the crossover operation. Common nodes between these two parents were determined. A random number of these were assigned to both the children symmetrically with a probability known as the crossover rate (70%). Unique nodes were randomly shuffled (with crossover rate) among the children. Mutation then randomly changed the network. New values replaced rate parameters according to the parameter mutation rate (10%). The parameter ranges used during mutation are shown in **Table 1**. In addition, non-existent edges could be duplicated (12%), and existing edges can be deleted (12%) with a defined probability.
2. Each child network was subsequently simulated to obtain the likelihood values using (6).
3. Networks that make up the next generation were determined by their scores. Here we used the deterministic crowding method, which started by calculating the similarity between the parent and child networks [45]. This method maintained diversity while improving the average likelihood value generation after generation. To calculate the distance metric, we used the following equation for Parent1 and Child1:

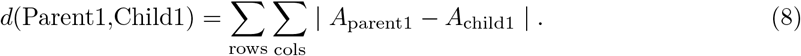

**Table 1.** Parameter ranges used for inference using genetic algorithms and inference algorithms.

| Parameter | Range |
| --- | --- |
| $k'$ | 0-1 |
| $K'$ | 0-15 $hr$ |
| $\gamma$ | 0-1 $hr^{-1}$ |
| $n$ | 1-3 |
| $D$ | 0-1000 $um^2/s$ |

Here *A* is the adjacency matrix of interaction between genes, with *A*_*ij*_ representing the regulation of gene *i* by gene *j* (1= activation, 0= none, −1= repression). Similar distance metrics were calculated for all parent and children pairs. Once we obtained the measure *d*, we used it to determine the parent that competes with a particular child for a spot in the next generation. This was determined by the likelihood value of each network (**Figure 2**). These steps were performed iteratively for many generations to converge to a population of networks.

**Figure 2.**
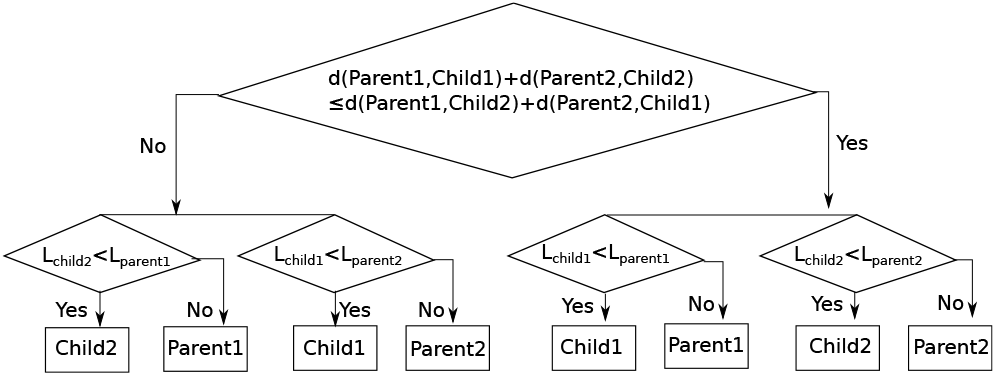
Description of the selection process in the genetic algorithm. First, the cross and related distances were compared starting with two parents and two children. Depending on the outcome, parent and child likelihood values are compared to give the four selected networks for the next generation. This method was adapted from [45].

A few modifications had to be made to improve the efficiency of this algorithm. The likelihood term for the 2D co-culture experiment was set to zero in Eqn (6) until the last 300 generations due to the high computational cost of simulating it. Note here that the 2D co-culture spatiotemporal data could contain signatures of autoregulatory feedback in terms of a spatial gradient of reporter protein expression, which follows that of secreted factors. Hence, after introducing spatiotemporal data in the last 300 generations, edge duplication was allowed in the autoregulatory feedback edges of genes. Also, for each of the first 500 generations, ten additional networks were inserted into the population with complementary networks to the existing networks to maintain diversity.

#### 2.4.2 Parameter inference

Once the directed graph representing the intercellular GRN was obtained, continuous rate parameters of each interaction (constituting the edge weights) were inferred. We used a different scoring function for parameter inference than the genetic algorithm for edge inference. Whereas scaling of spatial and temporal fluorescence between the minimum and maximum values as in (7) and truncating the time span improves inference of edge type, it alters spatiotemporal dynamics. However, normalizing the concentration of the reporter protein with respect to mean fluorescence preserves these dynamics and allows for inference of continuous rate parameters. The mean also averages out the measurement noise and reduces its impact on the normalized variable. The likelihood function employed in this case was given by:

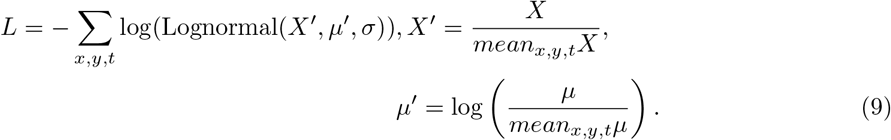

We assumed the likelihood function was based on a lognormal probability density function from experimentally observed fluorescence intensity distributions [46–49]. The lognormal function was defined such that the simulated value exp(*μ*′) corresponded to the median value of the measurements *X*′ and *σ* represented the standard deviation of the measurement noise of the corresponding normal distribution. Heuristic optimization algorithms like particle swarm in MATLAB with parallel processing enabled can be used to infer parameters with set upper and lower bounds on parameters shown in Table. 1. These are then fed as initial guesses into steepest-descent nonlinear programming solvers (e.g. fmincon in MATLAB) for local optimization. The optimized parameters values serve as initial estimates for subsequent algorithms, determining the uncertainty in the estimates given measurement noise.

#### 2.4.3 Uncertainty estimation in parameters given the noise in fluorescence reporters

To quantify parameter uncertainty given noisy gene expression data, we used the Bayesian method known as Markov Chain Monte Carlo (MCMC), specifically the Metropolis-Hastings algorithm with Gibbs sampling. Given prior estimates of parameters, the algorithm constructs a chain of parameter samples with each new iteration conditioned on the previous one. The resulting ensemble of sample values from many iterations made up the posterior distribution. This represented the joint probability distribution of parameter values in the model that explained the data given noise. The joint probability distribution may also indicate practical identifiability issues in the form of pairwise correlations between different parameters. This knowledge could be used to reduce the number of free parameters further.

A component of the MCMC sampling is the likelihood function, given in (9). The prior distributions may be centered around the parameters obtained from the inference methodology described in the previous section. The prior distributions were assumed to be log-normal, with standard deviations of 30% of the mean for the Hill coefficient *n*, 10% for *k*′, and 50% for other parameters.

After we set the likelihood function and the prior distributions, the MCMC algorithm progressed such that at each iteration, several values were proposed for a particular parameter. If *L* at the proposed point was greater than at the current point, the move was always accepted. Otherwise, it was only accepted with probability proportional to the ratio of *L* at the new point and the old point. Intuitively, it is visited less often to ensure that the long-term frequency of visits is proportional to its underlying probability. The parallelized version of this algorithm generated a probability distribution of *n*_*c*_ number of proposals sampled from the prior distribution where each point’s probability was proportional to the likelihood ratio with respect to the current point [50]. The computation of the likelihood function, the most expensive step, was carried out in parallel with *n*_*c*_ cores. Next, *n*_*c*_ new points were sampled from this distribution and one point at random was used for the next iteration to continue the chain.

To execute this algorithm, we assumed that the true parameters were inferred, and the prior distribution mean was centered around them. This would be possible with sufficiently large computational effort and time. We ran the MCMC for 10^5^ iterations, with *n*_*c*_ = 30 at each iteration leading to 3 *×* 10^6^ iterations. The objective was to obtain uninformative posterior distributions with a standard deviation at most half that of prior distributions. If the acceptance rate was lower than 1%, then the standard deviations were scaled down such that the prior distributions were accepted between 1% and20% of the time. If the prior distribution was informative with the mean of the posterior more than two standard deviations away from the prior mean, we shifted the prior distribution and reran the MCMC.

## 3 Results

### 3.1 *In silico* validation

We took three experimentally relevant cases to validate our methodology. We obtained the data set by using the lognormal function to simulate measurement noise and perturb the original *in silico* data with a standard deviation assumed to be 5% of the mean. Experimentally this value may be obtained using the radial average and the standard deviation of spatial data for the 2D co-culture experiment and replicates from the conditioned media experiment. The conditioned media experiment had a time interval of 0.3 hours and a total time of 72 hours. For the 2D spatiotemporal simulation we set the dimension of the geometry to be 40mm x 40mm discretized with 25 × 25 square elements. The 2D spatiotemporal data were acquired at intervals of 4.5 hours for a total of 45 hours. We set the maximum number of genes per cell as three, with one gene being a reporter gene, another the corresponding gene of interest which secretes an intercellular diffusible species. The third possible gene produces strictly intracellular regulatory proteins. This framework allows for at least 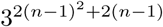 configurations of intercellular GRNs with *n* genes per cell, e.g. three genes/cell leads to 531441 configurations.

### 3.2 Case study 1: sender-receiver activation

We tested the inference on the sender-receiver activation using an established experimental model where the senders secrete the diffusible factor *Ahl* catalyzed by the *LuxI* gene and the receivers have plasmids with *Ahl* inducible *GFP* [51, 54]. The *LuxI* gene in the senders is activated by *Ahl*’s presence. This positive feedback loop mediated by *Ahl*, is one mechanism used for the quorum sensing observed in nature.

A system of equations described the time evolution of the species concentration as follows:

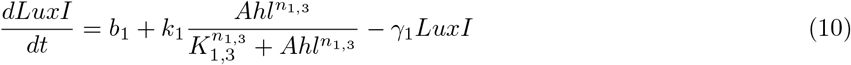

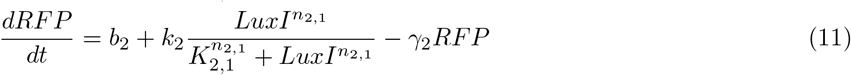

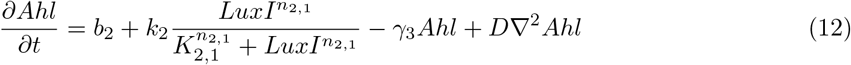

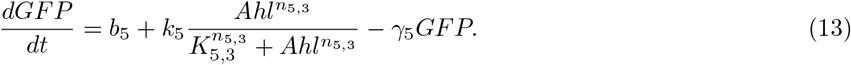

Using these equations and parameters given in Table. 2, we generated *in silico* reporter expression data corresponding to the 2D spatiotemporal experiment and the temporal conditioned media experiment (**Figure 3**). Note that *LuxI* is in quasi steady-state due to a high activation rate mediated by *Ahl* (*k*_1_). This means the level of *LuxI* ≫ *K*_2,1_, and consequently, reduced expression of reporter expression away from the center of the sender disk was not observed (**Figure 3B**). This caused the genetic algorithm to significantly reduce the inferred network (**Figure 4C**) compared to the true network (**Figure 4B**) and eliminated the positive feedback loop. The negative log-likelihood value was reduced with increasing generations of the genetic algorithm (**Figure 4D**).

**Table 2.** Parameters used for the sender-receiver activation system.

| Parameter | Value | Parameter | Value |
| --- | --- | --- | --- |
| $b_1$ | $10^{-3}$ nM/hr [51] | $\gamma_1$ | $0.024 \text{ hr}^{-1}$ [51] |
| $k_1$ | 25000 nM/hr [51] | $\gamma_2$ | $0.04 \text{ hr}^{-1}$ [52] |
| $b_2$ | 0 nM/hr [51] | $\gamma_3$ | $0.005545 \text{ hr}^{-1}$ [53] |
| $k_2$ | 0.06 nM/hr [51] | $\gamma_5$ | $0.04 \text{ hr}^{-1}$ [52] |
| $k_5$ | 0.06 nM/hr [51] | $n_{1,3}$ | 2 [51] |
| $K_{1,3}$ | 3.16 nM [51] | $n_{2,1} = n_{3,1}$ | 1 [51] |
| $K_{2,1}$ | 1 nM [51] | $n_{5,3}$ | 1.5 [54] |
| $K_{5,3}$ | 10 nM [54] | $D$ | $100 \text{ } \mu\text{m}^2/\text{s}$ [54] |

**Figure 3.**
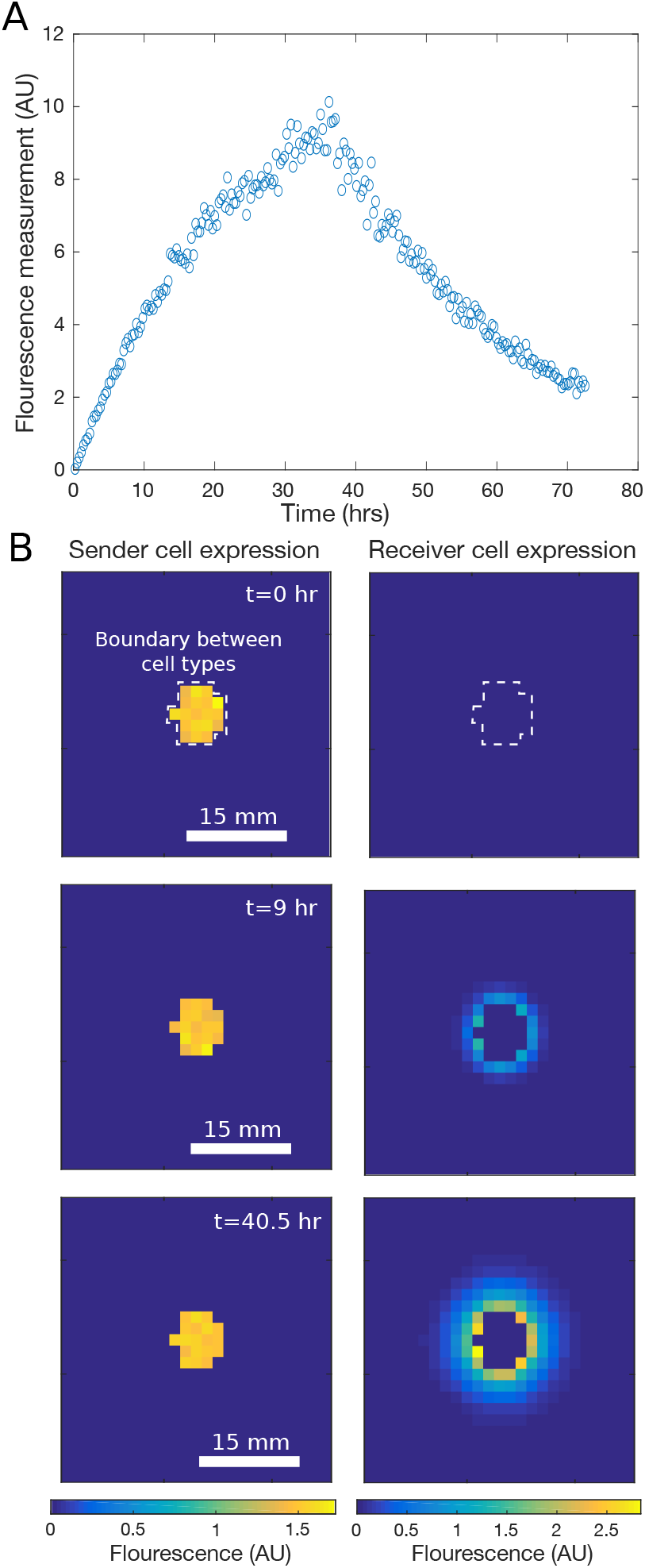
*In silico* data generated from sender-receiver activation model simulation. A: Temporal reporter fluorescence in receiver cells in a well-stirred volume induced at *t* = 0 by the introduction of a bolus of inducing factor and washed with conditioned media to remove it at *t* = 35*hrs*. B: 2D spatiotemporal reporter expression from sender and receiver cells in a co-culture experiment. The sender and receiver cells begin interaction at *t* = 0 and shown are three of ten time points used for inference. The control volumes were assigned cell types according to an arbitrary geometry (dotted white line being the boundary). Measurement noise was assumed to be lognormal with standard deviation as 5% of the mean.

**Figure 4.**
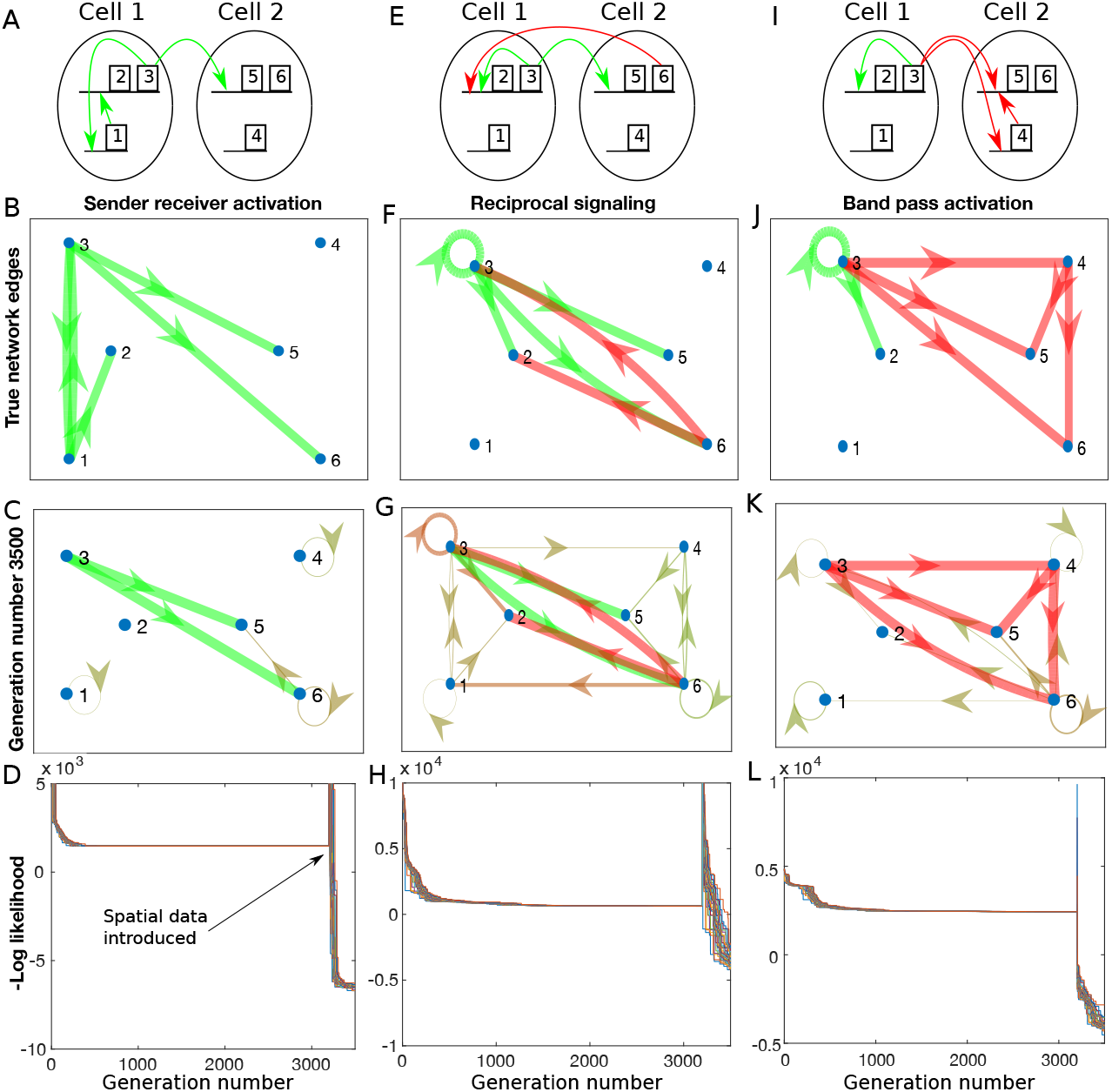
Network edge inference using genetic algorithm: A-D: Shows the true network used to generate *in silico* data for the sender-receiver case using a schematic (A), or graph representation (B). Genes 1, 2, 3, and 5 are *LuxI, RFP, Ahl* and *GFP*, respectively. *RFP* and *GFP* are reporters of the *Ahl* gene in senders and a gene of interest (gene 6) in receivers respectively. Red arrows represent repression and green represent activation. The GA converged to the inferred network shown in graph representation with uncertain edges given by off-colored thinner edges (C), while (D) is the likelihood function for each child in the population. Spatial data was introduced after generation no. 3200, thus, the likelihood value for generation *>* 3200 cannot be compared with that in the generations *≤* 3200. We stopped the GA when the network converged (even though likelihood values do not completely plateau) and can switch to a more efficient algorithm for edge parameter inference. E-H: Reciprocal signaling inference, where genes 2, 3, 5, and 6 are *GFP, A, RFP* and *B* respectively, I-L Sender receiver band-pass inference, where genes 2, 3, 4, 5 and 6 are *RFP, A, B, GFP* and a receiver gene of interest, respectively.

Identifiability analysis on the inferred network showed that the parameters 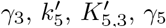, and *D* were uniquely identifiable from the inferred network assuming fluorescence measurement of the reporter. The parameters of the positive feedback loop (the second term in (10) and (11)) were absent in the inferred network. This means *k*_2_ = 0 in the inferred network. Note that the Hill coefficient *n*_5,3_ was assumed to be identifiable given the spatial gradient of *GFP* in the receiver cells. Relaxing the assumption of the availability of the absolute measurement of the reporter concentration means that the parameters must be scaled according to (5). Production parameters using (5) are 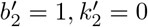and hence were not parameters to be inferred. The degradation rate of *RFP* was a parameter that was identifiable only when the absolute concentrations were available as 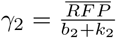where 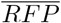 is the steady-state level of *RFP* and assuming *b*_2_ and *k*_2_ are known. As *RFP* concentration is not an excitable state variable, it remains at steady-state without any temporal dynamics, and thus *γ*_2_ cannot be inferred. Further, even though the scaled inducible *GFP* reporter gene expression rate was identifiable, we fixed it to its true value 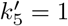prior to inference. This is because, in the unfixed case, the posterior distribution will asymptotically skew towards one and this value will be excluded from any *<* 100% confidence interval owing to one being the upper limit of the range of 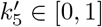.

The parameters of the resulting network were successfully estimated to be within 3% of their true values (**Figure 5A**). The highest deviations are for the degradation rate of *Ahl γ*_3_ and the scaled half-saturation constant of activation of *GFP* 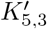, with deviations of opposite signs. The increased sensitivity of *GFP* to *Ahl* offset the high degradation rate of *Ahl* in the inferred network and gave a relatively unchanged likelihood value. This points to a practical identifiability issue where given measurement noise, the likelihood value is insensitive to parameters deviations in a small neighborhood around the true values.

**Figure 5.**
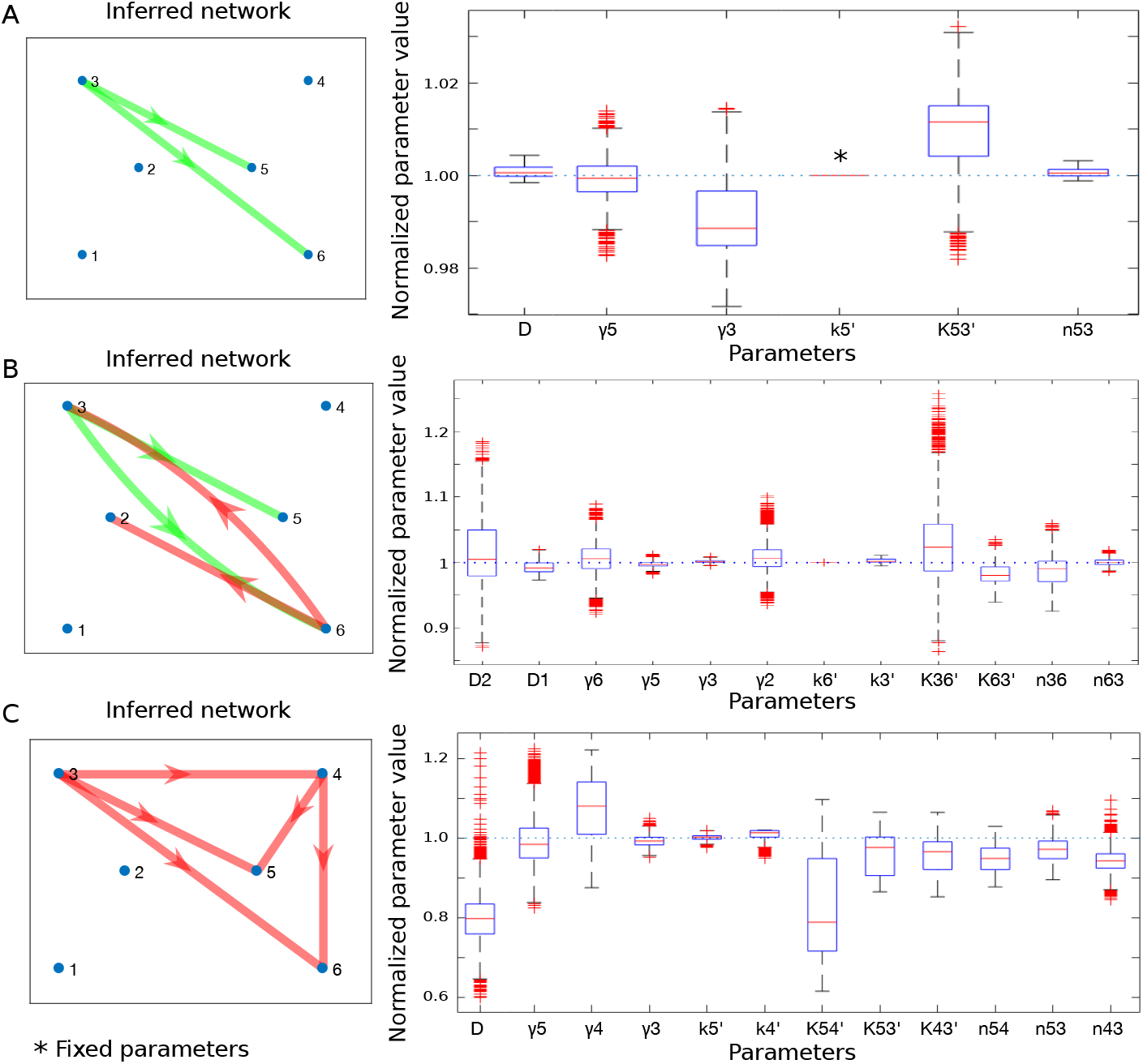
Network rate parameter estimation using MCMC. Each subfigure shows the uncertainty of the identifiable rate parameters (scaled) corresponding to the sender-receiver activation (A), reciprocal signaling (B), and bandpass expression (C) network inferred using the GA algorithm. The posterior parameter samples are shown as a ratio with respect to true scaled parameter values in a box plot format. Box edges are the 25^*th*^ and 75^*th*^ percentile of values, with the central red line being the median value, outliers represented by (+) markers, and the whiskers are the extreme data points excluding outliers. Outliers were defined as data points outside the 1.5*×* interquartile range about the box edges. Parameters with fixed values were at their biologically plausible range’s upper or lower limit.

### 3.3 Case study 2: reciprocal signaling

Broadly in multicellular systems, reciprocal signaling is a fundamental process used to guide morphogenesis [29, 55]. These systems have been synthesized in bacterial cells using plasmids to generate complex spatiotemporal patterns [56]. We choose a reciprocal signaling system with parameters shown in **Table 3**. The system of ordinary differential equations used was:

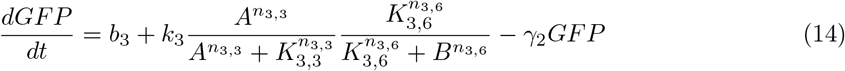

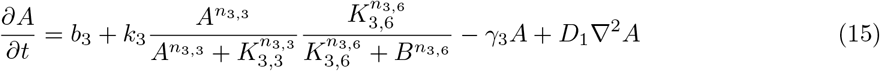

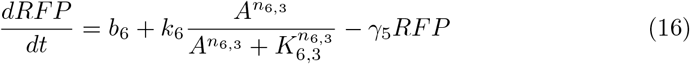

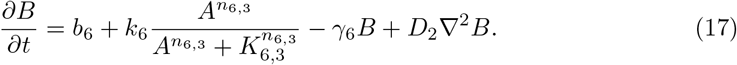

**Table 3.** Parameters used for the reciprocal signaling system.

| Parameter | Value | Parameter | Value |
| --- | --- | --- | --- |
| $b_3$ | 3 nM/hr | $\gamma_3$ | $0.4436 \text{ hr}^{-1}$ |
| $b_6$ | 0.8 nM/hr | $\gamma_5$ | $0.1663 \text{ hr}^{-1}$ |
| $k_3$ | 30 nM/hr | $\gamma_6$ | $0.1663 \text{ hr}^{-1}$ |
| $k_6$ | 40 nM/hr | $n_{3,3}$ | 2.5 |
| $K_{3,3}$ | 0.1 nM | $n_{3,6}$ | 1 |
| $K_{3,6}$ | 2.5 nM | $n_{6,3}$ | 2 |
| $K_{6,3}$ | 0.15 nM | $D_1$ | $100 \text{ } \mu\text{m}^2/\text{s}$ |
| $\gamma_2$ | $0.4436 \text{ hr}^{-1}$ | $D_2$ | $100 \text{ } \mu\text{m}^2/\text{s}$ |

**Table 4.** Parameters used for the band detect system.

| Parameter | Value | Parameter | Value |
| --- | --- | --- | --- |
| $b_3$ | 6 nM/hr | $K_{5,4}$ | 100 nM |
| $b_4$ | 1 nM/hr | $\gamma_3$ | $0.3 \text{ hr}^{-1}$ |
| $b_5$ | 0.8 nM/hr | $\gamma_4$ | $0.2 \text{ hr}^{-1}$ |
| $k_3$ | 60 nM/hr | $\gamma_5$ | $0.15 \text{ hr}^{-1}$ |
| $k_4$ | 50 nM/hr | $n_{3,3}$ | 1.5 |
| $k_5$ | 40 nM/hr | $n_{4,3}$ | 2 |
| $K_{3,3}$ | 20 nM | $n_{5,3}$ | 2 |
| $K_{4,3}$ | 20 nM | $n_{5,4}$ | 2 |
| $K_{5,3}$ | 20 nM | $D$ | $300 \text{ } \mu\text{m}^2/\text{s}$ |

Here the first cell type has an auto-regulatory feedback loop and secretes an activating factor, while the second cell type secretes a repressor in response.

Using the genetic algorithm, the likelihood value increased with increasing generation number, and the network converged to a simple activation-repression reciprocal signaling pattern (**Figure 4G**).

Similar to the sender-receiver case, the positive feedback loop in the cell type one was not inferred using the genetic algorithm. The repression from the second cell type could not be distinguished from the autoregulatory positive feedback in the spatial data.

Identifiability analysis of the inferred network, showed that all of the parameters in (14)-(17) re-scaled as in (5) were identifiable. The Hill coefficients were assumed to be equal to one for tractable identifiability analysis of the other parameters. We assumed that they would be identifiable from the gradient of the reporter expression and included them for inference. This assumption was validated, as the posterior distributions of all the parameters, including the Hill coefficients, were narrower than the priors. Additionally, all the inferred parameter confidence intervals included the true value (**Figure 5B**), with the upper and lower bounds being within 20% of their true value.

### 3.4 Case study 3: sender-receiver bandpass expression

The sender-receiver bandpass is the case where the sender secretes a factor, which auto-activates its own secretion. The receiver cells subsequently sense the extracellularly secreted factor, and only cells within a specific spatial area express fluorescence. This is possible by having a high detect and a low detect component regulating the reporter expression together in the receiver cells [57]. The intercellular GRN used is shown as a schematic (**Figures 4I**) and network graph (**Figures 4J**), with the parameters of the model set such that the 2D spatiotemporal data illustrates the band expression feature in receiver cells induced by the sender cells. The original system of differential equations was:

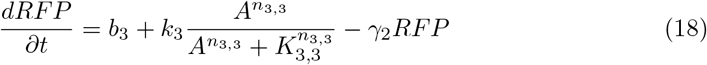

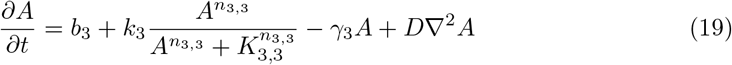

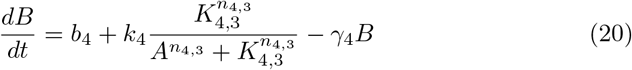

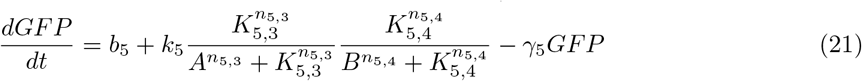

The genetic algorithm successfully inferred the incoherent feedforward circuit and the corresponding AND type regulation of GFP reporter expression in receiver cells (**Figure 4K**). The senders’ positive feedback loop was absent, similar to the previous cases. In some runs of the algorithm, the double repressive arm of the gene regulation in the receiver cells was substituted with double activation. This demonstrates that network edges may not be identifiable in some cases, and multiple networks may explain the data with similar likelihood values.

The identifiable parameters were 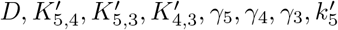 and 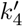. As there were no sender-reporter dynamics and *γ*_2_ represented the time scale of its response, STRIKE-GOLDD classifies *γ*_2_ as unidentifiable.

The inferred parameters deviated more than in the previous cases (**Figure 5C**). This is not surprising compared to the sender-receiver activation case due to higher dimensionality (12 vs 6 parameters). In comparison to the reciprocal signaling, the deviations were higher for two reasons. First, the inference was carried out using data from two reporter proteins vs only one in the bandpass case. Secondly, the inference of kinetic parameters of the intermediate protein (gene product 4) without a corresponding reporter using downstream reporter measurements (gene product 5) was an indirect task and was very sensitive to noise. Additionally, the significant underestimation of the diffusion constant was due to its correlations with parameters related to sensitivity, such as the half-saturation constants 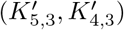 and Hill coefficient (*n*_4,3_) and their slight deviations from true values.

## 4 Discussion

This work used spatiotemporal data to infer network edges and parameters between genes of interest in two cell types. While we have limited the maximum number of potential unknown genes involved in the intercellular GRN, single-celled RNA-seq-based methods can be used before our inference methodology to obtain a plausible network between genes of interest [25, 58, 59]. This can serve as an initial seed network for the inference using the genetic algorithm we propose. Our inference methodology produces the smallest intercellular GRN network whose edges actively engage to create the observed spatiotemporal gene expression dynamics.

Agent-based tools like BSim [28, 60] enable the modelling of intercellular gene interactions with high resolution. The system of ODEs corresponding to the intracellular GRN in each cell are simulated in the system with spatial heterogeneity in extracellular diffusible factor concentration. However, given a large number of agents (cells) present in realistic systems, this modeling technique would be computationally infeasible for the task of parameter inference. Hence, we opted to model the intercellular interactions at the macroscale scale in these studies.

Prior to parameter inference, parameters need to be classified as structurally identifiable. We also developed a framework for using the STRIKE-GOLDD MATLAB package [44] to investigate parameter identifiability using spatiotemporal measurements of reporter proteins. Re-scaling parameters, and using the framework we obtained fully structurally identifiable models. Parameter estimates given measurement noise were then obtained using the Markov Chain Monte Carlo sampling.

The likelihood functions in this work were inspired by experimentally observed fluorescence distributions. However, in the absence of an *a priori* model of the measurement noise [61], repeated experiments at specific points in time and space could give the type of distribution and relationship between the mean and standard deviation [62]. For the 2D co-culture experiment, in lieu of multiple experiments, a single experiment with cell types organized in a radially symmetric geometry (e.g. petri dish) can be used to obtain repeated measurements and estimate the likelihood function.

The caveat of this methodology is that we do not provide a way to infer *a priori* whether parameters are practically identifiable. This issue arises when multiple sets of parameters generate similar fits to the data within the tolerance of measurement noise, despite the parameters being structurally identifiable [63]. This leads to some of the higher deviations from true values in specific parameters (**Figure 5**). The inference of network edges using the genetic algorithm also suffers from this issue wherein autoregulatory feedback loops are not inferred in networks. One remedy is to perform multiple replicates of experiments and include the data for fitting. This will increase the constraining power on the parameter space, increasing the likelihood of inference of a globally unique parameter set. For instance, in the case study of reciprocal signaling, a larger disc of sender cells is liable to show a more prominent sender reporter gradient following the spatial gradient of the diffusible factor. This would increase the likelihood of inference of the autoregulatory positive feedback loop. This issue will be systematically tackled in future work where an additional step will be incorporated to extend the scope beyond structural identifiability. An altered MCMC method [64, 65] or profile likelihood [63] have been used in previous biophysical models and are two candidates for this step.

Overall, this work tackles the problem of inference of intercellular gene interaction between different cell types. While intercellular inference using spatiotemporal gene expression has been carried out, it was restricted to interactions between cells of the same type [27, 28]. In our methodology we consider multiple cell types with potentially different intracellular GRNs interacting through morphogens. More importantly, unlike these previous studies, our inference methodology does not assume a concentration measurement of the diffusible factor. In addition, while identifiability of parameters is presumed in these studies, we propose parameter scalings and model identifiability tools to ensure the parameters in our models are uniquely identifiable. High throughput studies can connect a large number of genes across multiple cell types [25]. However, this requires resource intensive assays like *in situ* hybridization and scRNA-seq, and does not provide rate constants. While our methodology tackles low complexity GRNs with a low number of genes/cell, it gives kinetic rate constants using data from simple experiments. Finally, while we have considered *in vitro* experiments with assigned cell geometries, similar experiments with tissues cultures can also be used for inference, with the caveat that the reaction-diffusion control volume size may have to be reduced to describe the more complicated cell type boundaries.

## 5 Acknowledgements

The authors thank members of the Gleghorn and Zurakowski research groups for productive discussions and feedback on the draft manuscript.

